# “Identifying and characterizing scene representations relevant for categorization behavior”

**DOI:** 10.1101/2023.08.17.553708

**Authors:** Johannes J.D. Singer, Agnessa Karapetian, Martin N. Hebart, Radoslaw M. Cichy

## Abstract

Scene recognition is a core sensory capacity that enables humans to adaptively interact with their environment. Despite substantial progress in the understanding of the neural representations underlying scene recognition, the relevance of these representations for behavior given varying task demands remains unknown. To address this, we aimed to identify behaviorally relevant scene representations, to characterize them in terms of their underlying visual features, and to reveal how they vary across different tasks. We recorded fMRI data while human participants viewed scenes and linked brain responses to behavior in three tasks acquired in separate sessions: manmade/natural categorization, basic-level categorization, and fixation color discrimination. We found correlations between categorization response times and scene-specific brain responses, quantified as the distance to a hyperplane derived from a multivariate classifier. Across tasks, these effects were found in largely distinct parts of the ventral visual stream. This suggests that different scene representations are relevant for behavior depending on the task. Next, using deep neural networks as a proxy for visual feature representations, we found that early/intermediate layers mediated the relationship between scene representations and behavior for both categorization tasks, indicating a contribution of low-/mid-level visual features to these representations. Finally, we observed opposite patterns of brain-behavior correlations in the manmade/natural and the fixation task, indicating interference of representations with behavior for task demands that do not align with the content of representations. Together, these results reveal the spatial extent, content, and task-dependence of the visual representations that mediate behavior in complex scenes.

## 2. Introduction

Humans rapidly process scene information, allowing them to flexibly categorize and adaptively react to their immediate environment. Such highly efficient categorization relies crucially on the visual system, which extracts visual features from the environment and integrates them into increasingly complex representations through a series of hierarchically organized brain regions in the ventral visual stream (Epstein & Baker, 2019; Grill-Spector & Weiner, 2014; Op de Beeck et al., 2008). While this hierarchy of representations underlies successful categorization, the extent to which particular scene representations in the ventral visual stream are relevant for categorization behavior is poorly understood. Specifically, it remains unknown i) where in the brain scene representations relevant for behavior emerge, ii) what visual features these representations capture, and iii) to what degree the relevance of these representations for behavior varies given different task demands.

Concerning the first question, previous studies have used diverse methods to identify visual representations of simple and complex stimuli that are relevant for categorization behavior (DiCarlo & Maunsell, 2005; Majaj et al., 2015; Philiastides et al., 2006; Philiastides & Sajda, 2006). One such method particularly suited for complex real-world stimuli is the neural distance-to-bound approach (Ritchie & Carlson, 2016), which links visual representations in the brain to behavioral responses via the distance of brain responses from a hyperplane in a high-dimensional response space estimated by a multivariate classifier. Analogous to signal detection theory (Green & Swets, 1966), where distance from a criterion negatively correlates with reaction time, points close to the hyperplane indicate weak sensory evidence, leading to longer RTs, while points far from the hyperplane indicate strong sensory evidence, resulting in short RTs. Thus, given a negative relationship between neural distances and behavioral response times (RTs), the approach assumes that information in a given brain area is behaviorally relevant i.e. represented in a format that could potentially be read out into behavior (e.g. by an upstream area).

Using this approach, behaviorally relevant object representations have been identified in early visual as well as high-level object selective regions (Carlson et al., 2014; Grootswagers et al., 2018; Ritchie & Op de Beeck, 2019). A recent study has extended these insights to representations of complex scenes, demonstrating that scene representations relevant for manmade versus natural categorization behavior arise in a time window from 100-200 ms after stimulus onset (Karapetian et al., 2023). However, where in the brain such scene representations emerge remains unknown.

Concerning the second question, i.e. the visual features that behaviorally relevant representations capture, prior research has suggested that representations in scene-selective regions capture a variety of visual features, ranging from low to high level of complexity (MacEvoy & Epstein, 2011; Stansbury et al., 2013; Watson et al., 2014). However, for some basic distinctions such as categorizing scenes as manmade or natural, low-level visual features such as the spatial frequency or the color of a scene may be sufficient (Oliva & Torralba, 2001). This suggests that not all visual features that are captured by scene representations might be required for every scene categorization behavior and raises the question of what visual features underlie behaviorally relevant scene representations.

Thirdly, scenes can be categorized according to various criteria, such as manmade or natural, open or closed, or as belonging to a certain basic-level category (e.g. a beach, a highway etc.), with systematic differences in behavioral responses across tasks (Greene & Oliva, 2009; Kadar & Ben-Shahar, 2012; Loschky & Larson, 2010). While some of these differences might be accounted for by image-level properties (Sofer et al., 2015), they may also reflect more fundamental differences in the way neural representations are translated into behavior during these tasks. For tasks that require access to information not aligned with the content of scene representations, the represented information might even interfere with task performance (Greene & Fei-Fei, 2014; Reeder et al., 2015; Seidl-Rathkopf et al., 2015; Wyble et al., 2013). However, to what extent varying task demands influence the relationship between scene representations and behavior remains unclear.

Here, we identified behaviorally relevant scene representations in the brain, characterized them in terms of their underlying visual features, and investigated how they vary given different task demands. For this, we linked fMRI data from human participants viewing scene images to behavioral responses acquired in separate behavioral experiments for either a manmade/natural categorization task, a basic-level categorization task on the same scene images, or an orthogonal task on the fixation cross. To identify behaviorally relevant scene representations in the brain, we first localized scene category representations using multivariate decoding (Haynes & Rees, 2006) and then determined which of these representations are relevant for manmade/natural or basic-level categorization behavior by employing the neural distance-to-bound approach (Ritchie & Carlson, 2016). Next, to elucidate the nature of the behaviorally relevant representations, we determined what type of visual features, quantified as activations from different layers of deep neural networks, best explained these representations. Finally, to investigate how tasks that do not align with the content of scene representations impact the behavioral relevance of scene representations, we related scene representations to behavior in an orthogonal fixation task.

## 3. Materials and Methods

### 3.1. Participants

30 healthy adults with normal or corrected-to-normal vision participated in the fMRI study. All participants provided their written informed consent before taking part in the study and were compensated for their time. One participant was excluded from the analyses due to incidental findings consistent with a recognized neurological disorder, resulting in a final sample of 29 participants (mean age = 24.4, SD=3.7, 21 female, 8 male). The final sample size is comparable or larger than previous studies using decoding approaches for relating brain data to behavioral data (Carlson et al., 2014; Grootswagers et al., 2018; Karapetian et al., 2023; Ritchie & Op de Beeck, 2019). The study was approved by the ethics committee of Freie Universität Berlin and was conducted in accordance with the Declaration of Helsinki.

### 3.2. Experimental stimuli

We used 60 individual scene images from the validation set of the large-scale scene dataset Places365 (Zhou et al., 2018) (Fig. 1A). Half of the images depicted manmade scenes and the other half natural scenes. The images were further subdivided into 6 basic-level categories (beach, canyon, forest, apartment building, bedroom, highway), with 10 exemplars for each category. To standardize the size and aspect ratio of the stimuli, all images were center cropped and resized to 480×480 pixels.

**Figure 1.**
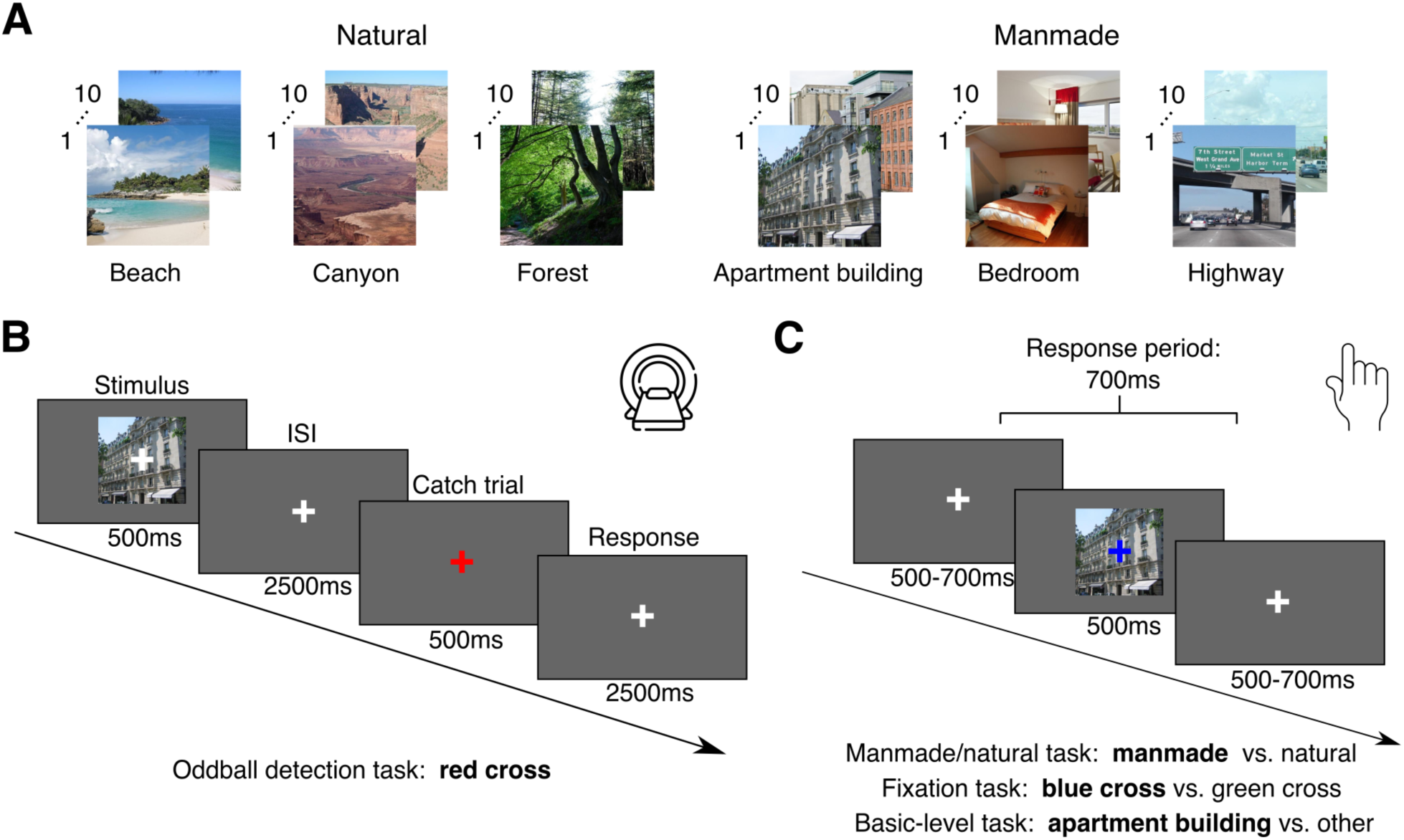
Stimulus set and experimental paradigm. **A) Stimulus set used in the experiment.** We used 60 scene images from the validation set of the Places365 dataset (Zhou et al., 2018). Half of the stimuli depicted manmade and the other half natural scenes and spanned 6 basic-level categories: beach, canyon, forest, apartment building, bedroom, highway. **B) fMRI paradigm.** In a given trial, a scene image was presented for 500ms overlaid with a white fixation cross, followed by an interstimulus interval (ISI) of 2500ms. In 20% percent of the trials the fixation cross turned red instead of the stimulus presentation and participants were instructed to press a button. **C) Behavioral paradigm.** Behavioral data was acquired with different sets of participants in either a previous experiment (Karapetian et al., 2023) or in an independent behavioral experiment with analogous trial structure but a different behavioral task. In a given trial, a scene image was presented for 500ms, overlaid with a blue or green fixation cross (only for the manmade/natural and fixation tasks), followed by the presentation of a white fixation cross for a variable time between 500-700ms. In the different experiments, participants were instructed at the beginning of a block to either report if a given scene image was a manmade or natural scene (manmade/natural task), if the color of the fixation cross was green or blue (fixation task), or if a given scene image belonged to a certain basic-level category of scenes or not (basic-level task).

### 3.3. Experimental design and procedure

#### 3.3.1. fMRI experimental paradigm

During the main fMRI experiment, participants were presented with individual scene images while fixating. Stimuli were presented for 500ms at 12 degrees of visual angle (width & height), overlaid with a central white fixation cross subtending 1 degree of visual angle (Fig. 1B). This was followed by an interstimulus interval of 2,500ms. In 20% of the trials, the fixation cross turned red instead of a stimulus presentation, and the participants were tasked to respond with a button press. Stimulus order was pseudo-randomized within a given run, avoiding immediate repetition of the same stimulus. Each participant completed either 8 or 10 runs, with each run lasting 7min 46.5s. In a given run each stimulus was presented twice, resulting in 16 or 20 stimulus repetitions in total for a given participant.

#### 3.3.2. Functional localizer task

To define regions of interest (ROIs), participants completed a functional localizer run at the beginning of the recording session. The localizer consisted of 15s blocks of objects, scrambled objects and scenes (not used in the main experiment) interleaved with 7.5s blocks of only the fixation cross on background as baseline. The images were displayed at a size of 12 degrees of visual angle, at the center of the screen for 400ms, followed by a 350ms presentation of the fixation cross. Participants were instructed to maintain fixation on the fixation cross and to press a button in case the same image was presented in two consecutive trials. In total, the localizer run included 8 blocks of each image type, resulting in a duration of 7min 22.5s. The order of the blocks was pseudo-randomized, avoiding immediate repetition of the same type of block.

### 3.4. fMRI acquisition, preprocessing and univariate analysis

#### 3.4.1. fMRI acquisition

We collected MRI data using a Siemens Magnetom Prisma Fit 3T system (Siemens Medical Solutions, Erlangen, Germany) with a 64-channel head coil. Structural scans were acquired using a standard T1-weighted sequence (TR=1.9s, TE=2.52ms, number of slices: 176, FOV=256mm, voxel size=1.0mm isotropic, flip angle=9°). Functional images were acquired using a sequence with partial brain coverage (TR=1s, TE=33.3ms, number of slices: 39, voxel size: 2.49×2.49mm, matrix size=82×82, FOV=204mm, flip angle=70°, slice thickness=2.5mm, multiband factor=3, acquisition order=interleaved, inter-slice gap=0.25mm). The acquisition volume fully covered the occipital and temporal lobes. Due to a technical update of the scanner the voxel size as well as the FOV was slightly changed for the sequence used in the localizer experiment for 20 out of the 30 participants (voxel size: 2.5×2.5mm, FOV=205mm).

#### 3.4.2. fMRI preprocessing

We preprocessed the fMRI data using SPM12 utilities (https://www._l.ion.ucl.ac.uk/spm/) and custom scripts in MATLAB R2021a (www.mathworks.com).

We realigned all functional images to the first image of each run, slice-time corrected them and co-registered them to the anatomical image. Further, based on the functional images and tissue probability maps for the white matter and cerebrospinal fluid, we estimated noise components using the aCompCor method (Behzadi et al., 2007) implemented in the TAPAS PhysIO toolbox (Kasper et al., 2017). Finally, we smoothed the functional images of the localizer run with a Gaussian kernel (FWHM=5). The functional images of the experimental runs were not smoothed.

#### 3.4.3. fMRI univariate analysis

We used a general linear model (GLM) to model the fMRI responses to each scene image in a given run. As the regressors of interest, we entered the onsets and durations of each of the 60 scene images, convolved with a hemodynamic response function (HRF). As nuisance regressors, we entered the noise components and the movement parameters as well as their first and second order derivatives. In order to account for task- and region-specific variability in the HRF (Polimeni & Lewis, 2021) we employed an HRF-fitting procedure as described in (Prince et al., 2022). For this, we repeated the GLM fitting 20 times, each time convolving all of the regressors of interest with a different HRF obtained from an open-source library of HRFs derived from the Natural Scenes Dataset (Allen et al., 2022). After fitting all the GLMs, we extracted the beta parameter estimates for the scene image regressors from the GLM with the HRF that had resulted in the minimum mean residual for a given voxel. Please note that this approach does not introduce any positive bias to multivariate decoding analyses, since it only focuses on maximizing the overall fit to the data without using any condition-specific information. This procedure resulted in 60 beta maps (one for each scene image) for each run and participant.

For the localizer experiment, we used a separate GLM to model the fMRI responses. Onsets and durations of the blocks of objects, scrambled objects and scenes defined regressors that were convolved with the canonical HRF. We only included movement parameters as nuisance regressors in this GLM. For localizing functionally defined brain areas, we computed three contrasts: scrambled > objects to localize early visual brain areas, objects > scrambled to localize object-selective cortex, and scenes > objects to localize scene-selective cortex. This yielded three *t*-maps for each participant.

#### 3.4.4. Region-of-interest (ROI) definition

As ROIs, we defined early visual cortex (EVC) i.e. V1, V2, and V3, as well as object-selective lateral occipital complex (LOC) and scene-selective parahippocampal cortex (PPA). For the definition of all ROIs we followed a two step procedure. First, we used masks based on a brain atlas with anatomical criteria for EVC (Glasser et al., 2016) and masks based on functional criteria for LOC and PPA (Julian et al., 2012). We transformed these masks into the individual subject space. Next, we computed the overlap between the subject-specific masks and the corresponding *t*-maps from the localizer experiment and only retained the overlapping voxels with *p*-values smaller than 0.0001. For EVC, we used the scrambled > objects *t*-map, for LOC we used the objects > scrambled *t-*map and for PPA we used the scenes > objects *t*-map. Finally, we excluded voxels that overlapped between any of the ROIs. This resulted in one EVC, LOC and PPA ROI mask for each subject.

### 3.5. Multivariate decoding of scene category information

To determine the amount of scene category information present in the fMRI response patterns we used multivariate decoding. For this, we trained and tested linear Support Vector Machine (SVM) classifiers (Chang & Lin, 2011) to distinguish whether a given fMRI response pattern belonged to a given scene category or not. We performed two types of decoding: manmade/natural decoding and basic-level decoding. For selecting train and test data for the classifiers, we used two different approaches: an ROI-based method targeting predefined regions and a spatially unbiased searchlight method for further specifying the spatial extent of local effects (Haynes et al., 2007; Kriegeskorte et al., 2006). We conducted all analyses separately for each subject and in the subject’s native anatomical space.

We formed pattern vectors based on the beta values from the voxels in a given ROI or searchlight. For this, we assigned all but four beta patterns for each scene image to the train set and the remaining four beta patterns to the test set. Please note that each beta pattern was based on data from a separate run, thereby avoiding potential false positives due to carry-over effects (Mumford et al., 2014). In order to improve the signal-to-noise ratio, for a given scene image we averaged betas from multiple runs into pseudo betas (Stehr et al., 2023). For the train set we averaged two betas into one pseudo beta and for the test set we averaged all four betas into one pseudo beta. Depending on whether participants finished 8 or 10 main experimental runs, this resulted in either 2 or 3 pseudo betas per scene image for the train set and one pseudo beta for the test set. For the manmade/natural decoding we used data for all of the images for training and testing the classifier. For the basic-level decoding, we sampled data for 10 target images belonging to the given scene category (e.g. apartment building) and 5 distractor images for each of the other two categories (i.e. bedroom, highway) within the same superordinate category (i.e. manmade) of the given target category, in order to balance the amount of positive and negative examples in the train set.

To increase the robustness of the results, we repeated the splitting of the data into train and test sets, sampling of target/distractor categories for the basic-level decoding, and the pseudo beta averaging 100 times while randomly shuffling the order of the betas. The resulting decoding accuracies were averaged across repetitions.

For the ROI-based method we iterated this procedure across ROIs and for the searchlight-based method across searchlights. This resulted in one decoding accuracy for manmade/natural decoding and 6 decoding accuracies for basic-level decoding (one for each target category) for every ROI and one searchlight decoding map for every subject. Decoding accuracies and decoding accuracy maps for basic-level decoding were averaged across target categories. For later group-level statistical analyses, we normalized the searchlight decoding maps to the MNI template brain.

### 3.6. Behavioral data

In order to identify behaviorally relevant scene representations, we linked the neural data recorded in the present study to behavioral data from three different tasks. Behavioral data for the manmade/natural categorization and fixation tasks was recorded in a previous study (Karapetian et al., 2023), while the data for the basic-level categorization task was recorded in an additional experiment with an independent set of 32 participants. One of these participants was excluded due to not finishing the experiment (final sample *N*=31, mean age=26.1, *SD*= 5.42, 24 female, 7 male).

In the manmade/natural categorization and fixation task experiment, 30 participants were presented with the same scene images as used in the fMRI study and performed either a manmade/natural categorization task on the stimuli or an orthogonal color discrimination task on the fixation cross (i.e. fixation task) while EEG was recorded. The experiment consisted of 20 blocks, 10 per task, and included at least 30 trials per scene image per block. In each trial, a stimulus was presented for 500ms overlaid with a green or blue (randomly assigned) fixation cross, followed by a presentation of a white fixation cross for a variable time window between 500 to 700ms. Participants were instructed at the beginning of each block to either report if the presented stimulus was a manmade or a natural scene or to report the color of the fixation cross, as accurately and as quickly as possible.

In the basic-level categorization experiment, participants were presented with the same scene images as in the experiments mentioned above and were instructed to indicate with a button press if the present image belonged to a given basic-level scene category (e.g. apartment building) or not. The trial structure was equivalent to the other behavioral experiment but fixation cross color change trials were not included. At the beginning of each block, participants were informed which basic-level scene category to categorize and were given example images (distinct from the experimental stimuli) for that given category. In a given block only the 10 exemplar images of the given scene category and randomly sampled distractor images from the same superordinate category (manmade/natural) were presented. The experiment consisted of 24 blocks, 4 per scene category, and included 24 trials per image.

For all three behavioral experiments separately, we first averaged the response time (RT) data from the correctly answered trials for each subject and then averaged RTs across subjects to obtain the mean RT for each scene image and each task. On average, for a given subject, 23.2 (*SD* = 6.0) trials were included for each scene for the manmade/natural task, 26.0 (*SD* = 1.46) for the fixation task, and 20.8 (*SD* = 3.79) for the basic-level task. This resulted in one mean RT for each scene image and each task.

### 3.7. Distance-to-bound analysis

We used the neural distance-to-bound approach (Carlson et al., 2014; Ritchie et al., 2015; Ritchie & Carlson, 2016) to determine if scene information represented in fMRI response patterns is behaviorally relevant for a given task (Fig. 2A). The neural distance-to-bound approach links the information in brain patterns to behavior by predicting a relationship between RTs and distances of individual brain responses to a criterion in the high-dimensional neural response space. The concept of a criterion is based on signal detection theory (Green & Swets, 1966) and can be formulated in high-dimensional spaces as a hyperplane that is estimated when using multivariate decoding. The approach assumes a negative relationship between distances of individual brain response patterns to the hyperplane and RTs: points close to the hyperplane have weak sensory evidence and are difficult to categorize, leading to longer RTs. Vice versa, points far from the hyperplane have strong sensory evidence and can be easily categorized, resulting in short RTs. If this predicted relationship holds true for observed brain response patterns and behavioral responses, then it is assumed that information represented in these brain patterns is relevant for behavior.

**Figure 2.**
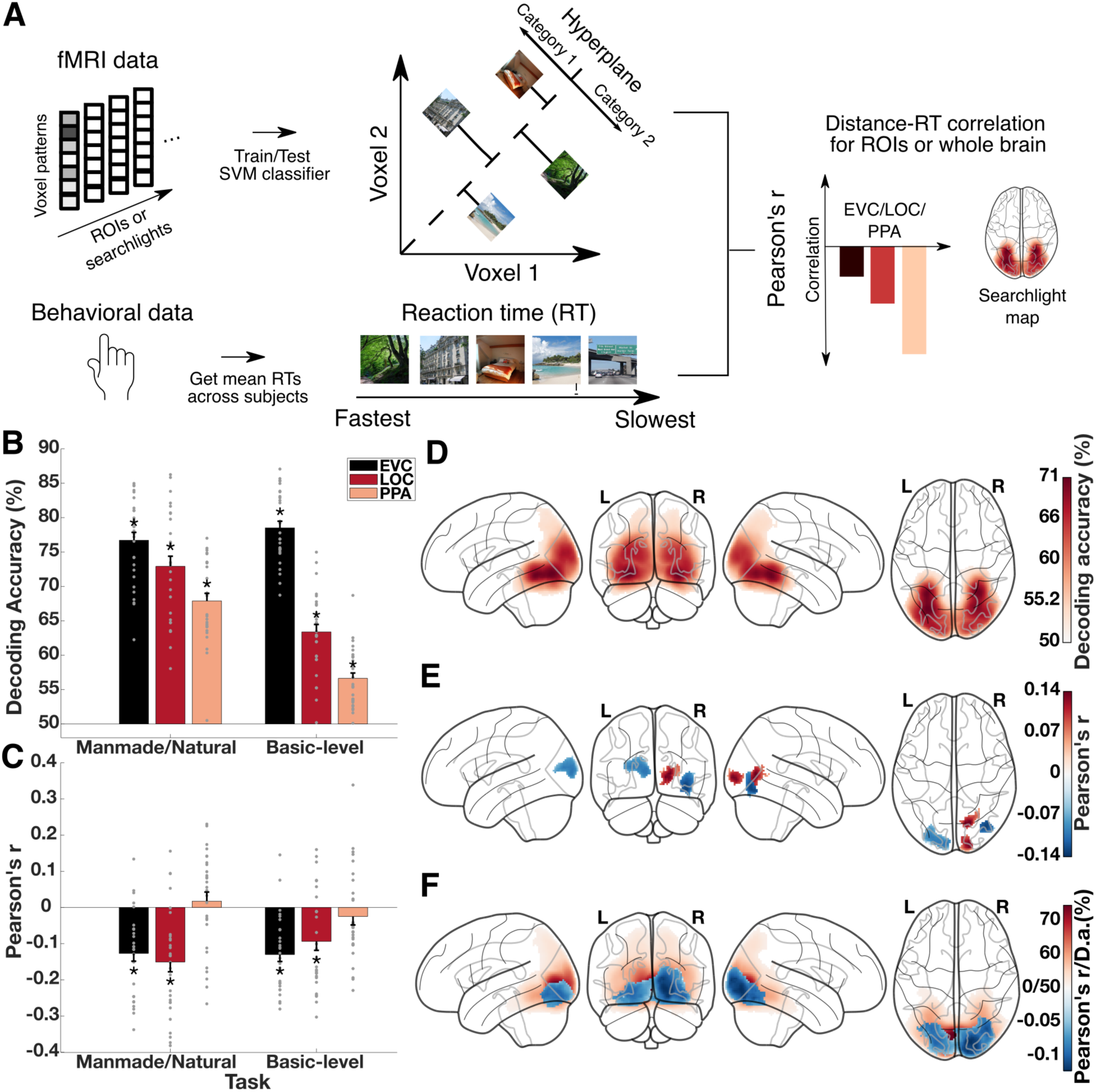
Scene category representations and behaviorally relevant scene representations in visual cortex. **A) Neural distance-to-bound approach for identifying behaviorally relevant scene representations.** For each subject, we derived neural distances from the fMRI response patterns by training SVM classifiers on part of the fMRI data and obtaining scene-specific distances from the hyperplane of the classifier for the left-out fMRI data for manmade/natural and basic-level decoding separately. Next, we obtained mean RTs (in a manmade/natural categorization task or a basic-level categorization task) across participants for each scene image and linked these RTs to the neural distances separately for each task using Pearson’s correlation. We iterated this procedure over ROIs or searchlights, resulting in ROI-specific correlation values or searchlight correlation maps. Negative correlations between neural distances and RTs at a specific location in the brain indicate that the representations at this location are relevant for behavior. **B) Scene category decoding in EVC, LOC and PPA.** Basic-level category as well as manmade/natural category could be decoded with accuracies significantly above chance in EVC, LOC and PPA. **C) Distance-RT correlations in EVC, LOC and PPA.** There were negative correlations between behavioral RTs and neural distances for both manmade/natural and basic-level categorization in EVC and LOC, but not PPA. Grey points indicate data points for individual subjects. Error bars depict the standard error of the mean across participants. Stars above or below the bars indicate significant results (*p*<.05, FDR-corrected). **D) Manmade/natural decoding across the visual cortex.** Searchlight manmade/natural decoding revealed significant decoding accuracies that were most pronounced in posterior and lateral parts of occipital cortex, with decreasing accuracies towards anterior parts of ventral-temporal cortex and posterior-parietal cortex. **E) Distance-RT correlations for manmade/natural categorization across the visual cortex.** Iterating the distance-RT correlation for manmade/natural categorization across searchlights showed negative correlations that were strongest at the border between occipital and ventral-temporal cortex as well as at the border between occipital and posterior parietal cortex. There were additional significant positive correlations which were strongest in the right occipital cortex. **C) Basic-level decoding and distance-RT correlations for basic-level scene categorization across the visual cortex.** For basic-level scene categorization, there were significant decoding accuracies across the whole ventral and dorsal stream with a peak in occipital cortex. Negative distance-RT correlations were found in posterior and lateral parts of occipital cortex.

To test the predicted relationship between neural distances to the hyperplane and RTs, we obtained distances for every scene image using the hyperplanes estimated with the same decoding procedure as described above. For manmade/natural decoding these distances were all obtained from the same decoder, while for basic-level decoding these values were obtained from 6 different decoders (one for each target category) and concatenated subsequently. We iterated this procedure over ROIs and searchlights, resulting in a vector with 60 values (one for each scene image) for each ROI, and searchlight. Finally, we correlated the vectors of distances with the vector of mean RTs for each ROI and searchlight using Pearson’s correlation. This yielded distance-RT correlations for each ROI, searchlight and subject.

### 3.8. Model-based distance-to-bound analysis

To examine what type of visual features best explains behaviorally relevant scene representations in the brain given different tasks, we used the neural distance-to-bound approach in combination with deep neural network (DNN) modeling and commonality analysis (Mood, 1971; Reichwein Zientek & Thompson, 2006). The basic rationale (Fig. 3A-C) involved first extracting activations from different DNN architectures and layers as an approximation of visual feature representations at different levels of complexity (Bankson et al., 2018; Groen et al., 2018; Reddy et al., 2021; Xie et al., 2020). The assumption that these activations approximate a gradient of feature complexity is based on demonstrations of a hierarchical correspondence between representations in DNNs and the human brain (Cichy et al., 2016; Güçlü & Gerven, 2015). Next, in order to link neural network activations, brain response patterns and behavioral RTs, we derived distances to the hyperplane based on the neural network activations for the manmade/natural and basic-level task separately. Finally, to determine which model activations accounted for behaviorally relevant scene representations, we estimated the shared variance between model distances, neural distances and RTs from different tasks using commonality analysis.

**Figure 3.**
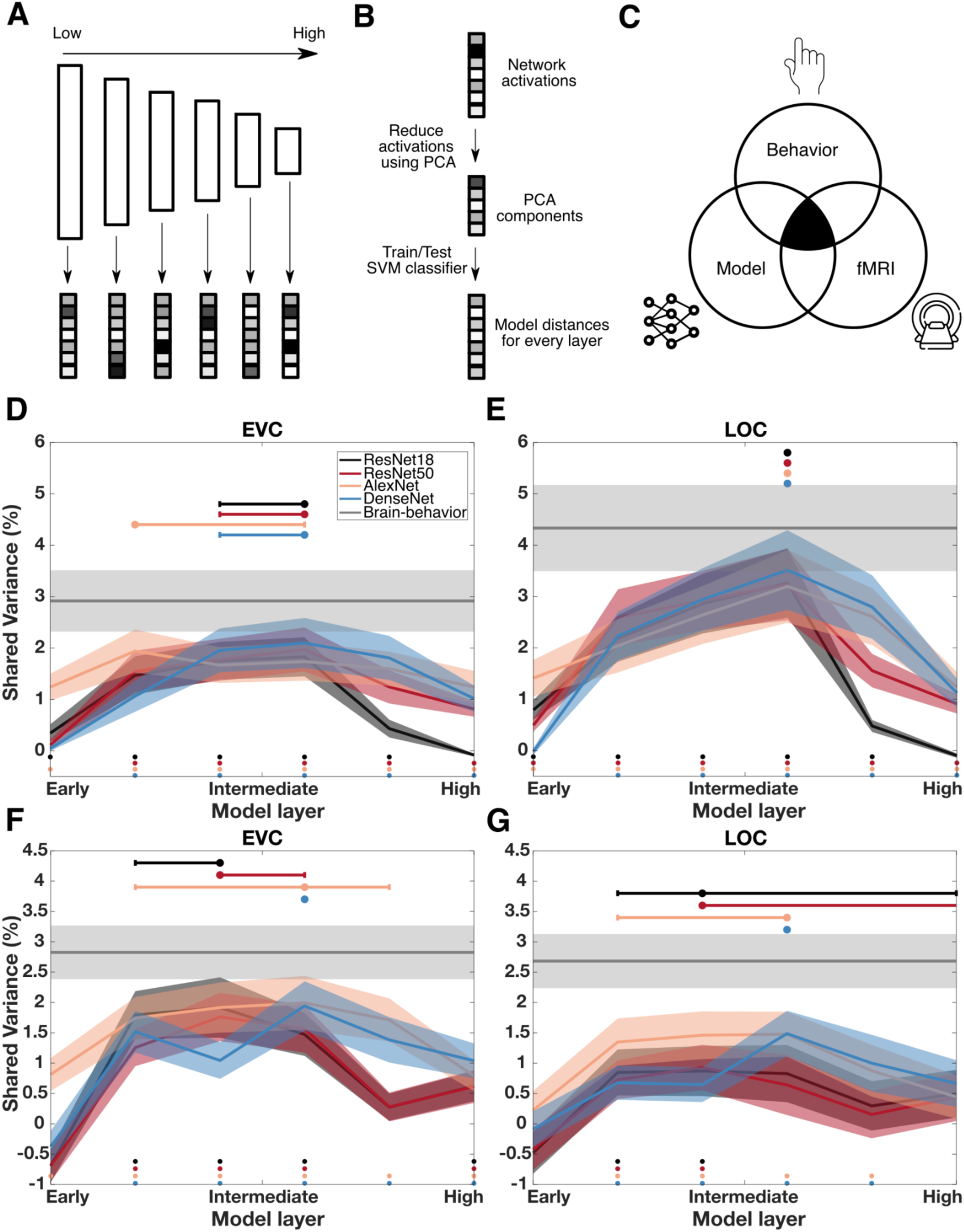
Visual features underlying behaviorally relevant scene representations. **A) Extraction of activations from various deep neural network layers.** As a proxy for visual feature representations, we extracted activations for scene images from the validation set of Places365 as well as for our experimental stimuli from various DNN architectures and layers. **B) Deriving scene-specific distances from neural network activations.** For linking the network activations to distances based on fMRI data and behavioral RTs we first reduced the activations using PCA. Next, for every layer and network separately, we trained SVM classifiers on either a manmade/natural or a basic-level scene classification task using the network activations and then tested the classifiers on the activations for our experimental stimuli. This yielded distances from the hyperplane for each of our experimental stimuli and every layer and network. **C) Commonality analysis approach.** To quantify how well model distances explain the shared variance between distances based on fMRI data and behavioral RTs, we assessed the shared variance between neural distances, model distances and behavioral RTs using commonality analysis for the manmade/natural and basic-level task separately. **D) Shared variance for manmade/natural categorization in EVC.** We found significant positive R^2^ values in all of the layers and networks except for the first layer in ResNet18, ResNet50 and DenseNet161 and the last layer in ResNet18. R^2^ values peaked in early/intermediate layers for all networks. **E) Shared variance for manmade/natural categorization in LOC.** R^2^ values were significant in all networks and layers except for the first layer in DenseNet161 and the last layer in ResNet18. For all networks R^2^ values peaked in intermediate layers. **F) Shared variance for basic-level categorization in EVC.** We found significant R^2^ values in layers 2-4 and 6 for ResNet18 and ResNet50, in all layers for AlexNet, and in layers 2-6 for DenseNet. R^2^ values peaked in early to intermediate layers for all networks. **F) Shared variance for basic-level categorization in LOC.** We found significant R^2^ values in layers 2-3 in ResNet18 and ResNet50, and in layers 2-5 in AlexNet and Densenet. R^2^ values peaked in early to intermediate layers for AlexNet and DenseNet. For ResNet18 and ResNet50 peaks were highly variable. Colored dots below the lines indicate significant layers. Shaded areas represent the SEM across participants. Horizontal error bars depict the 95% confidence intervals of the peak layer index. No horizontal error bar for a given layer indicates that the 95% confidence interval included only the value of the peak layer index. The gray line depicts the shared variance between brain distances and reaction times which corresponds to the upper limit for the shared variance between brain, models and behavior.

In detail, as models we used the ResNet-50, ResNet-18 (He et al., 2015), AlexNet (Krizhevsky et al., 2012) and DenseNet161 (Huang et al., 2018) architectures, pre-trained on the Places365 dataset (Zhou et al., 2018) (retrieved from https://github.com/CSAILVision/places365). We chose to examine different DNN architectures to ensure that a given pattern of results is not idiosyncratic to a given architecture but can be generalized to a given hierarchical level regardless of the specific architecture. For the manmade/natural task we extracted activations for 1,200 images from the validation set of Places365 (Zhou et al., 2018) as well as for our experimental stimuli. The Places365 images were sampled from 80 categories (half manmade, half natural), including the six categories from our stimulus set, and contained 15 images per category. For the basic-level task we extracted activations for a different set of images from the validation set of Places365 including 100 images for each of the 6 basic-level scene categories used in the experiment (i.e. 600 images in total) as well as for our experimental stimuli. For the extraction we focussed on a selection of layers including all pooling layers and the last fully connected layer for AlexNet, the output of all residual blocks and the last fully connected layer for the ResNets, as well as the first pooling layer, the output of all the DenseBlocks and the last fully connected layer for DenseNet161. For the manmade/natural task, we reduced the network activations for every layer to a dimensionality of 1,000 by using PCA on the activations for the 1,200 images from the validation set of Places365 (except for the fully connected layers that already had a dimensionality of <1,000). For the basic-level task, the dimensionality of the activations was reduced to 200 because of a lower amount of training samples than in manmade/natural classification. For both types of classification, we applied the estimated parameters to the activations for the train images as well as our main experimental stimuli.

Next, we trained and evaluated SVM classifiers separately for the manmade/natural and the basic-level task. For manmade/natural classification, we used the reduced activations for the 1,200 Places365 validation images for training for every layer and network separately, then tested the trained SVM classifiers on the reduced activations for our 60 experimental stimuli, and finally derived a distance to the hyperplane for each scene image. This resulted in 60 distances for each layer and network. For basic-level classification, we trained and evaluated a classifier for each of the 6 basic-level scene categories separately. We used the reduced activations of the 100 images for a given target category (e.g. apartment building) and from 50 randomly sampled images from both of the distractor categories (e.g. bedroom, highway) within the same the superordinate category (e.g. manmade) for training. We then tested the classifiers on the reduced activations from the 10 experimental images from the given target category and derived a distance to the hyperplane for each scene image, for every layer and network separately. To increase the robustness of the resulting distances, we repeated the sampling of target and distractor images 100 times and averaged the results subsequently. Finally, we concatenated the distances for the 10 test images of each target category, resulting in 60 distances for each layer and network.

Using commonality analysis, we finally determined the common variance between the network distances, neural distances and behavioral RTs for each task separately. In commonality analysis, the common variance that can be explained in a given outcome variable by two predictor variables is defined as the amount of variance explained by both predictors in the outcome variable minus the unique contribution of each of the predictors. In simplified form this term can be written as: C(AB)=R^2^_y.A_+R^2^_y.B_-R^2^_y.AB_, where R^2^ is the explained variance in a multiple regression model with the mean RTs as outcome variable (y) and either neural distances (A), network distances (B) or both (AB) as predictor variables. We fitted the corresponding multiple regression models and computed the commonality based on the R^2^ values, resulting in shared variance estimates for each network, layer, ROI, and subject.

### 3.9. Statistical analyses

For statistical testing we used non-parametric sign permutation tests at the group-level (Nichols & Holmes, 2002). We obtained null distributions for a statistic (decoding accuracies, distance-RT correlations) by randomly permuting the sign of the results at the participant level 10,000 times. Next, we obtained *p*-values for the observed data by comparing their statistic to that of the null distribution. We used one-sided tests for decoding accuracies and R^2^ values, as well as two-sided tests for distance-RT correlations and differences between decoding accuracies.

To correct for multiple comparisons, we used two different approaches. In the case of only a limited number of tests (i.e., < 10) such as multiple ROIs or neural network layers, we used the Benjamini-Hochberg FDR-correction without dependency (Benjamini & Hochberg, 1995). When applying a large number of tests such as for testing across searchlights (i.e., ∼ 100,000), we used a cluster-based correction (Maris & Oostenveld, 2007). For this, we first thresholded the *p*-values from the non-parametric sign permutation tests at *p*<0.001. Then we clustered the thresholded *p*-values by spatial adjacency and computed the maximum cluster size for each permutation. Next, we determined the *p*-value for each cluster in the observed data by comparing the cluster size of a given cluster to the maximum cluster size distribution. Finally, we thresholded the cluster *p*-values at *p*<0.05.

To compute 95% confidence intervals for the hierarchical level, i.e. the layer index where there was the peak R^2^ value obtained by the commonality analysis, we used bootstrapping. First, we took 100,000 random samples with replacement from the participant-specific R^2^ values. We computed the mean over participants for each bootstrap sample and detected the index of the layer with the peak R^2^ value across network layers. Finally, we used the 2.5% and 97.5% percentiles of the bootstrap distribution as the lower and upper bound of confidence intervals.

### 3.10. Data and code availability

The raw fMRI data is available in BIDS format on OpenNeuro (https://openneuro.org/datasets/ds004693). The beta maps obtained from the GLM, the behavioral data, the distances derived from the DNNs, as well as all first-level and group-level results are available via OSF (https://osf.io/y8tx2/). All code used for the first-level and group-level analyses in this study is provided via Github (https://github.com/Singerjohannes/visdecmak).

## 4. Results

### 4.1. Largely distinct representations in visual cortex are negatively correlated to RTs for different categorization tasks

First, in order to identify scene category presentations that could potentially be relevant for categorization behavior, we determined where information about scene category is present in the brain using multivariate decoding. For this, we trained SVM classifiers on the fMRI data to predict either if a given brain activity pattern belonged to a manmade or a natural scene or if the scene belonged to one of six basic-level scene categories (Fig. 1A) and tested the classifier on left-out data. We performed this analysis across three key regions of interest: early visual cortex (EVC), lateral occipital complex (LOC) and parahippocampal place area (PPA). Additionally, we employed a spatially-unbiased searchlight procedure (Haynes & Rees, 2006; Kriegeskorte et al., 2006) to uncover scene category representations beyond predefined ROIs. We performed significance testing using sign-permutation tests for all results. For a small number of multiple comparisons (<10, i.e. across ROIs, DNN layers) we applied an FDR-correction (Benjamini & Hochberg, 1995) and for multiple comparisons across searchlights we applied a cluster-based correction (Maris & Oostenveld, 2007).

For manmade/natural decoding as well as for basic-level decoding we found accuracies significantly above chance in all ROIs (*p*<0.001, Fig. 2B), suggesting the presence of scene category representations in these regions. This result was as expected from these regions’ central role in processing complex visual stimuli (Epstein & Baker, 2019; Grill-Spector & Weiner, 2014; Op de Beeck et al., 2008). Searchlight decoding revealed that manmade/natural decoding as well as basic-level decoding was significantly above chance (p<0.05) throughout the ventral and dorsal visual stream. Manmade/natural decoding was highest in posterior and lateral parts of the occipital cortex and decreased towards anterior parts of cortex (Fig. 2D), while basic-level decoding was strongest in posterior parts of the occipital cortex and decreased similarly towards anterior parts of cortex (Fig. 2F). Together, these results suggest a widespread presence of scene category representations as candidates for behaviorally relevant representations along both the ventral and dorsal stream (Walther et al., 2009, 2011).

Having identified scene category representations in the brain, we sought to determine to what extent these representations are relevant for different scene categorization tasks by using the distance-to-bound approach (Ritchie & Carlson, 2016, Fig. 2A). We first obtained mean RTs for the manmade/natural task and for the basic-level task across participants for each scene image. Then, we derived neural distances for each scene image from the SVM classifiers trained on the fMRI response patterns. We correlated these neural distances with behavioral RTs across the 60 scene images separately for each task and repeated this procedure across ROIs and searchlights.

For manmade/natural categorization, we found negative distance-RT correlations in EVC and LOC (both *p*<0.001, Fig. 2C) but not in PPA (*p*=0.488), suggesting that scene representations in EVC and LOC are relevant for manmade/natural categorization behavior, without positive evidence for a role of PPA. For basic-level scene categorization, we found negative distance-RT correlations in EVC and LOC (both *p*<.002, Fig. 2C), but not in PPA (both *p*=.304). This indicates that scene representations in EVC and LOC are relevant for basic-level scene categorization behavior, without positive evidence for PPA, and suggests that representations in similar brain regions contain information relevant for different scene categorization tasks.

Searchlight analysis further revealed significant negative distance-RT correlations (*p*<0.05, Fig. 2E) for manmade/natural categorization at the border between occipital and ventral temporal cortex and between occipital and posterior parietal cortex, but not in parahippocampal cortex. For basic-level scene categorization, negative distance-RT correlations were found in posterior and lateral parts of occipital cortex (*p*<0.05, Fig. 2F). To further investigate if the voxels with negative distance-RT correlations were distinct or overlapping between tasks, we calculated the overlap between the significance maps for the manmade/natural and basic-level scene task and quantified the overlap in percent of overall significant voxels for a given task. This revealed that only 2.21% of the significant voxels for the basic-level scene task and 13.01% of significant voxels for the manmade/natural task overlapped with the significant voxels of the other task, respectively. In contrast to the ROI results, this suggests that, while there is a partial overlap of behaviorally relevant representations for manmade/natural and basic-level scene categorization, the majority of representations for the two tasks are spatially distinct.

Surprisingly, we also found significant distance-RT correlations that were positive for manmade/natural categorization (*p*<0.05), which were confined to the right occipital cortex only. A positive correlation between neural distances and RTs violates the predictions of the neural distance-to-bound approach and suggests that a scene representation with a strong category signal leads to a slow RT in the task and vice versa. This implies interference between scene representations in the occipital cortex and behavior in the manmade/natural task.

Taken together, these results suggest that while there is a widespread presence of scene representations that are potentially relevant for categorization behavior across tasks, partially overlapping but largely distinct subsets of these representations in early visual and object-selective cortex, but not parahippocampal cortex, contain behaviorally relevant information depending on the task demands.

### 4.2. Features derived from early to intermediate neural network layers best explain behaviorally relevant scene representations in the visual cortex across tasks

While our findings so far suggest that largely distinct scene representations in the visual cortex are relevant for the different scene categorization tasks investigated, they leave open what types of visual features underlie these behaviorally relevant scene representations. We investigated this question in terms of feature complexity. As a proxy for low-to high complexity visual features, we used activations extracted from different layers of deep neural networks (for similar approaches see: (Bankson et al., 2018; Greene & Hansen, 2020; Groen et al., 2018; Reddy et al., 2021; Xie et al., 2020)) and asked to what extent these activations account for the link between scene representations and behavioral responses, separately for each task (for a visualization of the procedure, see Fig. 3A-C). We linked network activations to RTs and fMRI data using the neural distance-to-bound approach (Ritchie & Carlson, 2016) and determined which layer’s activations best explain the shared variance between RTs and fMRI data using commonality analysis (Mood, 1971; Reichwein Zientek & Thompson, 2006). We focussed on EVC and LOC since we found significant distance-RT correlations only in these regions. We applied right-tailed sign-permutation tests, testing for positive R^2^ values. We discuss the results ordered by task and then by region.

For manmade/natural categorization, we found significant R^2^ values in EVC for most of the networks and layers (all *p*<0.030, Fig. 3D) except for the first layer in ResNet50 and DenseNet161 and the last layer in ResNet18 (all *p*>0.168). In LOC we found significant R^2^ values for most networks and layers (all *p*<0.001, Fig. 3E) except for the first layer in DenseNet161 and the last layer in ResNet18 (both *p*>0.642).

For basic-level scene categorization, we found significant R^2^ values in EVC for layers 2-4 and 6 in ResNet18 and ResNet50, all layers in AlexNet, and layers 2-6 in DenseNet (all *p*<.028, Fig. 3F). In LOC we found significant R^2^ values in layer 2 and 3 for ResNet18 and ResNet50, and in layers 2-5 for AlexNet and DenseNet (all *p*<.039, Fig. 3G).

Together, these results demonstrate that for manmade/natural categorization, visual features from most hierarchical levels, excluding very early and late stages, and for basic-level scene categorization, visual features mostly from early to intermediate layers account for parts of the variance shared between brain and behavior.

Next, we determined which visual features explain the shared variance most strongly between brain and behavior by determining the layers with the highest shared variance. We use the following convention for reporting statistics: peak layer index [lower, upper] 95% (bootstrapped) confidence interval.

For manmade/natural categorization, we found that the shared variance in EVC peaked in early to intermediate layers for all networks (Fig. 3D, ResNet18 = 4 [3, 4], ResNet50 = 4 [3, 4], AlexNet = 2 [2, 4], DenseNet161 = 4 [3, 4]). In LOC, the shared variance peaked in intermediate layers for all networks (Fig. 3E, ResNet18 = 4 [4, 4], ResNet50 = 4 [4, 4], AlexNet = 4 [4, 4], DenseNet161 = 4 [4, 4]).

For basic-level scene categorization, we found that the shared variance in EVC peaked in early to intermediate layers in all networks (Fig. 3F, ResNet18 = 3 [2, 3], ResNet50 = 3 [3, 4], AlexNet = 4 [2, 5], DenseNet161 = 4 [4, 4]). In LOC, we found highly variable peaks for ResNet18 and ResNet50 (Fig. 3G, ResNet18 = 3 [2,6], ResNet50 = 3 [3,6]). This suggests that neither ResNet is a suitable model for explaining the shared variance between brain and behavior for basic-level categorization in LOC. For AlexNet and DenseNet values peaked in intermediate layers (AlexNet = 4 [2 4], DenseNet = 4 [4, 4]).

Together, these results suggest that behaviorally relevant scene representations for the two different categorization tasks are both best explained by low-to mid-level visual features.

### 4.3. Opposite brain-behavior correlations in a manmade/natural categorization task and an orthogonal fixation task

While we identified and characterized scene representations relevant for different scene categorization tasks, their relation to behavior might yet differ again for tasks that are not aligned with the represented scene information. Previous research showed that viewing scenes while performing an orthogonal task can impair performance (Greene & Fei-Fei, 2014; Reeder et al., 2015; Seidl-Rathkopf et al., 2015; Wyble et al., 2013). However, to what extent scene representations interfere with behavior in an orthogonal task remains unknown. To investigate this, we determined the behavioral relevance of scene representations for the orthogonal task of reporting the color of the fixation cross.

In the experiment investigating manmade/natural categorization, participants viewed scenes and colored fixation crosses simultaneously, while performing categorization and fixation cross color discrimination in alternating blocks. This suggests the hypothesis that the content of scene representations interacted with performance in the fixation task, which would be evident in a significant relationship between scene representations and fixation task RTs. Please note that participants in the fMRI experiment were neither engaged in the fixation task nor were they presented with different fixation cross colors during the presentation of the images. Thus, evidence supporting the above hypotheses would indicate that processing scenes, even when not task-relevant, engages representations that can interfere with performance in orthogonal tasks such as fixation cross color discrimination. Analogous to the procedures outlined above for identifying behaviorally relevant scene representations, here we correlated scene-specific distances derived from the manmade/natural decoders to RTs from the fixation task.

In contrast to the negative correlations for the manmade/natural task, we found positive correlations between neural distances and fixation task RTs in EVC and LOC (Fig. 4A, both *p*<0.003), but not in PPA (*p*=0.073). Searchlight analysis further revealed positive distance-RT correlations that were most pronounced at the border between occipital and ventral-temporal cortex (Fig. 4B, *p*<0.05). That is, scene representations with a strong category signal were found to be associated with slow responses in the fixation task and vice versa for scene representations with a weak category signal and speeded responses. In addition, we asked if the representations that showed a positive correlation with behavior in the fixation task were the same or different from the representations that exhibited a negative correlation with behavior in the manmade/natural categorization task. By computing the overlap between the voxels with significant negative correlations with RTs in the manmade/natural categorization and fixation tasks, we found only a small overlap of 9.48%. This indicates that scene representations relevant for the manmade/natural categorization and fixation task were largely distinct. Together, these findings suggest that scene representations at the border between occipital and ventral-temporal cortex, which are largely distinct from the behaviorally relevant representations for manmade/natural categorization, interfere with behavior in the fixation task.

**Figure 4.**
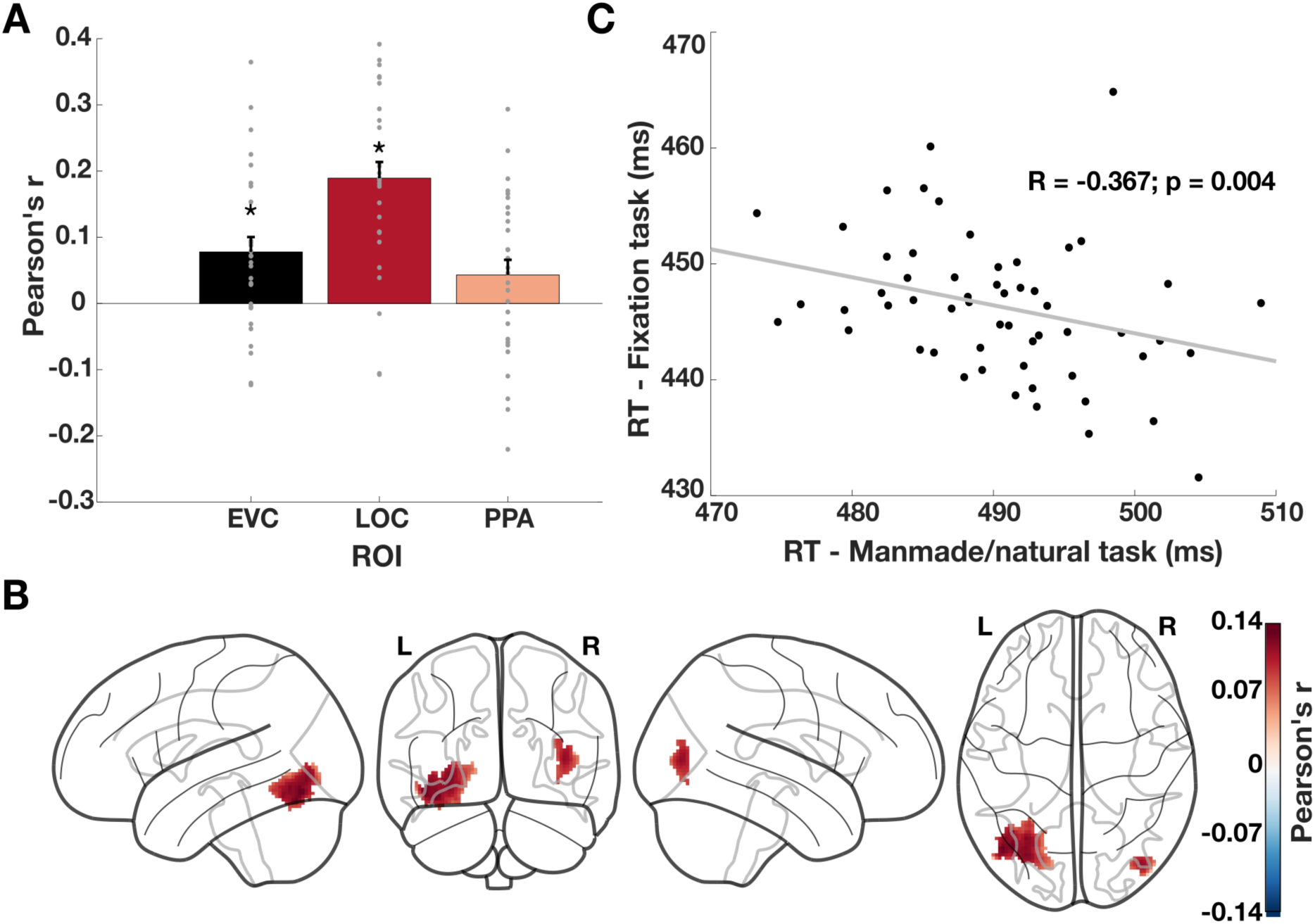
Behavioral relevance of scene representations for an orthogonal fixation task. **A) Correlations between RTs in the fixation task and neural distances in EVC, LOC and PPA.** We found significant positive correlations in EVC, LOC but not PPA. Grey points indicate data points for individual subjects. Error bars depict the SEM across participants. Stars above the bars indicate significant results. **B) Correlations between RTs in the fixation task and neural distances in the visual cortex.** We found positive correlations that were strongest at the border between occipital and ventral-temporal cortex. **C) Correlations between mean RTs for the manmade/natural and the fixation task.** We found a negative correlation between RTs in the fixation and the manmade/natural task.

A possible explanation for this interference might be that scene representations that evoke a strong category signal in the brain take away processing resources from the fixation task, thereby slowing the RT. Based on this explanation, we expected to observe a similar relationship between the RTs in the manmade/natural categorization task and the fixation task, namely that scenes that are solved faster in the manmade/natural categorization task lead to slower RTs in the fixation task and vice versa. To test this, we correlated the RTs from the manmade/natural categorization task with the RTs from the fixation task. We found a negative correlation between the RTs of the two tasks (*r*=-0.367, *p*=0.004; Fig. 4C), indicating that scene images that are solved fast in the manmade/natural task are associated with long RTs when presented during the fixation task and vice versa. This suggests that scene processing interferes with performance in the fixation task, corroborating the interference effect between scene representations and behavior in the fixation task.

In sum, these results provide evidence that a subset of scene representations in the visual cortex are relevant for behavior even in tasks beyond scene categorization. Yet, the relevance of scene representations for behavior differed for the fixation task and the scene categorization tasks. While scene representations facilitate categorization behavior, they interfere with behavior in the fixation task. This further corroborates the notion that the task demands critically affect the behavioral relevance of scene representations.

## 5. Discussion

In the present study we identified and characterized behaviorally relevant scene representations as well as their dependence on the task by relating fMRI responses to behavioral RTs in different tasks using the neural distance-to-bound approach (Ritchie & Carlson, 2016). The study yielded three key findings. First, while we could decode both manmade/natural as well as basic-level scene categories along both the ventral and dorsal stream, neural distances were negatively correlated to categorization RTs in largely distinct regions of the ventral visual stream for the two tasks. This suggests that despite a widespread and overlapping presence of scene category representations, depending on the task demands, mostly distinct subsets of these representations are behaviorally relevant. Second, distances derived from early to intermediate layers of deep neural networks best explained the shared variance between brain and behavior for both tasks, suggesting that low to mid-level visual features best account for behaviorally relevant scene representations for these tasks. Finally, we observed opposing patterns of correlation between neural distances and RTs for the fixation task and the manmade/natural task. While for manmade/natural RTs there was a negative correlation, suggesting facilitation of behavior, for fixation RTs we found a positive correlation, suggesting interference with behavior. This indicates that scene representations can interfere with behavior when there is a misalignment between these representations and the task demands. Together, these results elucidate the relationship between neural representations of scenes and behavioral performance by demonstrating how specific visual features and the task context mediate this relationship.

### 5.1. Largely distinct behaviorally relevant scene representations in visual cortex for different categorization tasks

By employing the neural distance-to-bound approach (Ritchie & Carlson, 2016), we identified partially overlapping but largely distinct scene representations relevant for manmade/natural and basic-level scene categorization behavior. These representations were localized at the border between occipital and ventral-temporal cortex for manmade/natural categorization and in posterior and lateral parts of occipital cortex for basic-level categorization, but interestingly not in parahippocampal cortex. These findings align with object recognition studies (Carlson et al., 2014; Grootswagers et al., 2018; Ritchie & Op de Beeck, 2019) showing behaviorally relevant representations in both early and high-level visual cortex and with studies suggesting that representations in different brain areas are flexibly accessed for different tasks (Birman & Gardner, 2019; Kang & Maunsell, 2020). Thus, our findings challenge the view that information for categorizing natural images is only read out from high-level visual cortex (Majaj et al., 2015) and suggest that representations from both early and high-level visual cortex might be flexibly read out in perceptual decision-making (Contier et al., 2024; Jagadeesh & Gardner, 2021).

Our findings complement and extend a recent characterization of behaviorally relevant scene representations over time (Karapetian et al., 2023) by spatially localizing these representations in the brain and by extending them to different scene categorization tasks. The presence of behaviorally relevant scene representations in LOC, but not PPA, is in line with studies emphasizing the role of LOC in scene recognition (Linsley & MacEvoy, 2014; MacEvoy & Epstein, 2011; Stansbury et al., 2013). However, the presence of behaviorally relevant representations in EVC and the absence of evidence of behaviorally relevant representations in PPA for any scene categorization task in our data conflicts with the pivotal role of PPA in scene recognition (Aguirre et al., 1998; Epstein & Kanwisher, 1998) and with findings of behaviorally relevant representations in PPA, but not EVC (Groen et al., 2018; King et al., 2019; Walther et al., 2009, 2011). One potential explanation for this discrepancy might be the information participants relied on for performing the tasks. Given the behaviorally relevant representations in LOC, which is associated with object representations, it is likely that participants relied on object information for the categorization tasks rather than other information such as spatial layout, which is more strongly associated with PPA (Park et al., 2011). This is in line with research suggesting that PPA primarily processes the spatial aspects of a scene rather than categorical divisions (Kravitz et al., 2011). Thus, in tasks emphasizing spatial aspects of a scene, PPA might be behaviorally relevant, while in tasks prioritizing other types of visual information, other regions might become behaviorally relevant. Another related explanation is the amount of processing time available to participants in our experiments. Previous experiments used very short image presentations (<50ms) followed by a mask, effectively limiting the depth of processing of the scene (Walther et al., 2009, 2011). This might constrain subjects to rely on more global features such as the layout of the scene for the task in contrast to more fine-grained information which is processed later (Bar et al., 2006; Hegdé, 2008; Schyns & Oliva, 1994; Sugase et al., 1999). In sum, rather than contradicting the pivotal role of PPA in scene processing, our findings suggest that other areas involved in processing scenes such as EVC and LOC might also represent behaviorally relevant information depending on the perceptually available information and the task demands.

Surprisingly, we found a positive correlation between neural distances in the right occipital cortex and RTs in the manmade/natural task. These findings are not captured by the original formulation of the neural distance-to-bound approach (Ritchie & Carlson, 2016), which assumes a negative relationship between neural distances and RTs, where large distances are associated with fast RTs and vice versa. Instead, we observed a case of the opposite pattern: large distances were associated with slow RTs and vice versa, suggesting interference between scene representations and behavior in the manmade/natural task. This interference is hard to reconcile with the role of the occipital cortex in visual processing. However, these positive correlations might be spurious and influenced by a bias in the classifier’s hyperplane towards a specific category (e.g. manmade, natural). Such biases in the distance-RT correlations towards one category of a given category division (e.g. animate over inanimate) have been reported previously (Carlson et al., 2014; Grootswagers et al., 2017, 2018; Karapetian et al., 2023; Ritchie et al., 2015). Fully understanding this phenomenon requires simulations of different data regimes in combination with an in-depth geometrical analysis of the estimated hyperplane and its relationship to individual data points, which is a promising avenue for future studies.

### 5.2. Low-to mid-level visual features best explain behaviorally relevant scene representations in the visual cortex across tasks

We found that low-to mid-level visual features best accounted for the shared variance between neural distances and RTs for both scene categorization tasks. These results align with findings highlighting the importance of low-level visual features such as color, pooled contrast, or spatial frequency (Groen et al., 2012, 2013; Oliva & Schyns, 2000; Oliva & Torralba, 2001) as well as mid-level visual features such as curvature or texture (Renninger & Malik, 2004; Walther & Shen, 2014) for scene categorization. However, our findings also diverge from previous studies, which showed that high-level conceptual features best explain variance in behavioral similarity judgments for scenes and objects (Greene & Hansen, 2020; King et al., 2019). One potential reason for this divergence is that similarity judgments might be based on different visual features than categorization RTs. While categorization RTs might depend on more perceptual information of low-to intermediate complexity (Eberhardt et al., 2016), judging the similarity of scenes might involve high-level features related to the semantics of a scene. Additionally, our findings challenge a body of research that has taken differences in RTs between manmade/natural and basic-level categorization as evidence for participants’ stronger reliance on global, rather low-level visual features for manmade/natural than for basic-level scene categorization (Kadar & Ben-Shahar, 2012; Loschky & Larson, 2010; Oliva & Torralba, 2001, 2006). Our results did not reveal differences in the type of visual features best explaining behaviorally relevant representations for these tasks. Such differences in visual feature use might be especially apparent under conditions of short presentation times and backward masking, where the amount of processing time biases humans to rely on the most rapidly available type of features. Given longer presentation times, as used in our experiments, participants might leverage other visual information in similar ways across tasks. Future studies might contrast different characterizations of behavior in response to scenes and their relationship to brain data with respect to the available processing time for a better understanding of the relevance of distinct types of visual features for various behavioral goals.

### 5.3. Interference of scene representations with behavior in orthogonal fixation task

We found opposing patterns of correlation between neural distances and RTs in the manmade/natural task and the fixation task. In the manmade/natural task, strong category signals were associated with fast RTs and vice versa, suggesting a facilitative relationship between scene representations and behavior. In contrast, for the fixation task, strong category signals were associated with slow RTs and vice versa. This suggests interference between scene representations and behavior in an orthogonal task. This interference could be due to automatic processing of the content of a scene (Greene & Fei-Fei, 2014) which might have interfered with the representation of the fixation cross color. Alternatively, attention might have been differentially captured by the scenes and diverted away from the fixation cross, thereby impairing performance in the fixation task (Reeder et al., 2015; Seidl-Rathkopf et al., 2015; Wyble et al., 2013). While our findings cannot distinguish between these alternatives, they highlight the importance of scene recognition as a core cognitive process which cannot be easily suppressed. Further, they corroborate that the behavioral relevance of scene representations critically depends on the task demands.

### 5.4. Limitations

Several experimental factors potentially limit the generalizability of our findings. First, our selected stimuli and tasks only represent a small subset of all possible tasks and naturalistic stimuli that could be used to investigate the link between scene representations and categorization behavior. The specific combination of task and stimulus set influences the representations and types of visual features that are relevant for the given behavioral responses, thus limiting our results to these particular choices. We believe that focusing on ecologically relevant tasks such as manmade/natural and basic-level scene categorization, using naturalistic stimuli that span a range of common scene categories, is a valuable step towards understanding the relationship between scene representations and behavior. However, a comprehensive understanding of this relationship necessitates large-scale neuroimaging datasets (Allen et al., 2022; Gifford et al., 2022; Hebart et al., 2023) in combination with a broad sampling of different behavioral tasks, which is an exciting future direction.

Additionally, our choice of task in the fMRI experiment might have limited the emergence of behaviorally relevant representations. Participants performed a change detection task on the fixation cross in the fMRI experiment which differed from both categorization tasks or the fixation task in the behavioral experiments and for which the scene images were not relevant. Even though previous studies have shown that performing a task on the experimental images is not necessary for emergence of behaviorally relevant visual representations in the occipito-temporal cortex (Carlson et al., 2014; Grootswagers et al., 2018), particularly representations in parietal and frontal brain regions are affected by the task (Bracci et al., 2017; Hebart et al., 2018; Vaziri-Pashkam & Xu, 2017). Thus, engaging participants in the same task in the fMRI and behavioral experiment could have expanded the detectable behaviorally relevant representations.

### 5.5. Conclusion

Together, our findings reveal the spatial extent of the visual representations underlying categorization behavior for real-world scenes, identify low-to mid-level visual features as the main contributor to these behaviorally relevant representations and suggest that the behavioral relevance of scene representations critically depends on the task context. These results contribute to the understanding of the neural mechanisms and visual features enabling adaptive perceptual decisions in complex real-world environments.

## Funding information

This work was supported by a Max Planck Research Group grant (M.TN.A.NEPF0009) of the Max Planck Society awarded to MNH, a European Research Council grant (ERC-StG-2021-101039712) awarded to MNH, the German Research Council grants (CI241/1-1, CI241/3-1, CI241/7-1) awarded to RMC, and a European Research Council grant (ERC-StG-2018-803370) awarded to RMC.

## Conflict of interest statement

The authors declare no competing financial interests.

## Acknowledgements

We thank Marleen Haupt, Alessandro Gifford and Tony Carricarte for comments on the manuscript. Computing resources were provided by the high-performance computing facilities at ZEDAT, Freie Universität Berlin. Some of the figures used in this paper have been designed using images from Flaticon.com

## Author contributions

**Johannes Singer:** Conceptualization, Investigation, Data Curation, Formal analysis, Software, Visualization, Writing - original draft; **Agnessa Karapetian:** Conceptualization, Data Curation, Investigation, Writing - reviewing & editing; **Martin Hebart:** Supervision, writing - reviewing & editing; **Radoslaw Cichy:** Conceptualization, Supervision, Funding acquisition, Writing - reviewing & editing

## Notes

### Competing Interest Statement

The authors have declared no competing interest.

### Summary of Updates

Major revision of the paper; includes new behavioral data with another scene categorization task and additional analyses for these new data

